# Ultrastructural analysis of intracellular membrane and microtubule behavior during mitosis of *Drosophila* S2 cells

**DOI:** 10.1101/232199

**Authors:** Anton Strunov, Lidiya V. Boldyreva, Evgeniya N. Andreyeva, Gera A. Pavlova, Julia V. Popova, Alena V. Razuvaeva, Alina F. Anders, Fioranna Renda, Alexey V. Pindyurin, Maurizio Gatti, Elena Kiseleva

**Author notes:** Correspondence:Anton Strunov, Maurizio Gatti.

## Abstract

S2 cells are one of the most widely used *Drosophila melanogaster* cell lines for molecular dissection of mitosis using RNA interference (RNAi). However, a detailed and complete description of S2 cell mitosis at the ultrastructural level is still missing. Here, we analyzed by transmission electron microscopy (TEM) a random sample of 144 cells undergoing mitosis, focusing on intracellular membrane and microtubule (MT) behavior. This unbiased approach allowed us to discover that S2 cells exhibit a characteristic behavior of intracellular membranes, involving the formation of a quadruple nuclear membrane in early prometaphase and its disassembly during late prometaphase. After nuclear envelope disassembly, the mitotic apparatus becomes encased by a discontinuous network of ER membranes that associate with mitochondria preventing their diffusion into the spindle area. We also observed a peculiar metaphase spindle organization. We found that kinetochores with attached k-fibers are almost invariably associated with lateral MT bundles that can be either interpolar bundles or k-fibers connected to a different kinetochore. This spindle organization is likely to favor chromosome alignment at metaphase and subsequent segregation during anaphase. In summary, we describe several previously unknown features of membrane and microtubule organization during S2 cell mitosis. The genetic determinants of these mitotic features of can now be investigated using an RNAi-based approach, which is particularly easy and efficient in S2 cells

## Introduction

Schneider 2 cells, commonly called S2 cells, are one of the most widely used *Drosophila melanogaster* cell lines. S2 cells were established from a primary culture of late stage embryos and are likely to be derived from a phagocytic hematopoietic cell lineage (Schneider, 1972; Echalier, 1997). Extensive work carried out in the past 15 years has shown that *Drosophila* S2 cells are an excellent system for molecular dissection of mitosis using RNA interference (RNAi). There are multiple advantages for using S2 cells in RNAi-based studies of mitosis. First, they have a very favorable cytology for spindle and chromosome visualization in both fixed and living cells (Goshima and Vale, 2003; Somma et al., 2008). Second, performing RNAi in S2 cells is very easy as it can be carried out by treating cultures with large dsRNAs (of 400-800 bp), which are incorporated by the cells and processed in small interfering RNAs (siRNAs) that efficiently degrade the target mRNA (Clemens et al, 2000; Meister and Tuschl, 2004). Additionally, simultaneous RNAi against multiple genes can be performed, allowing pathway dissection. Third, the *Drosophila* genome is fully sequenced and well-annotated and mitotic genes are highly evolutionary conserved among metazoans (Bier, 2005; see also FlyBase at http://flybase.org), allowing RNAi data on S2 cells to be extrapolated to human cells.

Over the past 15 years, hundreds of papers have been published exploiting RNAi to study the roles of individual *Drosophila* genes in S2 cell mitosis. RNAi in S2 cells was also used to analyze the functions of specific groups of genes such as those encoding for proteins involved in cytokinesis (Somma et al, 2002), microtubule-based motor proteins (Goshima and Vale, 2003), protein kinases (Bettencourt-Dias et al., 2004), and protein phosphatases (Chen et al., 2007). In addition, S2 cells were used to perform RNAi-based genome-wide screens for identification of genes involved in different aspects of cell division (Goshima et al., 2007; Somma et al., 2008) or in specific aspects of the process such as cytokinesis (Echard et al., 2004), centrosome assembly and behavior (Dobbelaere et al., 2008) or centrosome-independent spindle assembly (Moutinho-Pereira et al, 2013). Collectively, these studies identified many new mitotic genes and provided fundamental insight into the molecular mechanisms of cell division.

Despite the enormous work carried out on S2 cells mitosis, a detailed and complete description of the process at the ultrastructural level is still missing. Here, we analyzed S2 cell mitosis by transmission electron microscopy (TEM). We examined sections from a random sample of 144 cells undergoing mitotic division. This unbiased approach, allowed us to discover that S2 cells exhibit a peculiar intracellular membrane behavior, which allowed subdivision of prometaphase into four stages. Our observations also indicate that in late S2 cell prometaphase and metaphase, the spindle microtubules are mostly organized in bundles, which are either end-on attached to the kinetochores (k-fibers) or run from pole to pole (interpolar MTs). In addition, kinetochores with attached k-fibers almost invariably exhibit lateral associations with either k-fibers connected to different kinetochores or interpolar MT bundles, forming multi-bundle MT assemblies that are likely to aid chromosome alignment at metaphase and poleward movement in anaphase.

## Results and Discussion

To describe *Drosophila* mitosis at the ultrastructural level, actively proliferating *Drosophila* S2 cells were pelleted, fixed, embedded in resin and sectioned. Because cells within the pellet are randomly oriented, this procedure yielded sections along different planes of the mitotic apparatus. We examined only cells cut longitudinally or slightly obliquely with respect to the spindle axis (henceforth longitudinal sections), or cut transversally through the mitotic chromosomes (henceforth transverse sections); oblique cuts showing only part of the mitotic cell were discarded. For 67% of the cells, we analyzed single ultrathin sections (of approximately 70 nm). For the remaining cells, we performed serial sectioning and examined up to 15 sections per cell, mainly to obtain precise information on the intracellular membranes and kinetochore-MT relationships. For measuring cell structures we analyzed longitudinal and transverse 70 nm sections; in the case of serial sections, we considered only the central section of the cell.

Most of the cells in the pellet were in interphase, characterized by a rather homogeneous nucleoplasm and the presence of a prominent nucleolus; in the sections of these cells the nucleus was usually circular and surrounded by a double membrane (Figure S1A). We also observed cells that in addition to the nucleolus displayed several dense aggregations of chromatin and had the nucleus surrounded by an undulated double membrane; these cells might be in prophase or in a pre-prophase stage (Figure S1B, C).

We focused on 144 cells undergoing mitotic division: 71 prometaphases (49%), 37 metaphases (26%), 21 anaphases (15%) and 15 telophases (10%). In fixed S2 cells stained for tubulin and DNA and examined under a light microscope (n = 369), we found 40% prometaphases, 20% metaphases, 11% anaphases, and 29% telophases. Clearly, telophases are in excess compared to the TEM preparations. However, this is easily explainable, considering the difficulty in obtaining TEM sections with complete telophase figures including both presumptive daughter cells versus those that contain a single daughter cell, which were excluded from our analysis. If we do not consider telophases, the relative frequencies of prometaphases, metaphases and anaphases observed in TEM preparations (55%, 29% and 16%, respectively) and in light microscopy specimens (56%, 28% and 16%, respectively) are almost identical. Thus, at least for prometaphases, metaphases and anaphases, the TEM sample of mitotic cells we examined is representative of a population of actively proliferating S2 cells. We focused on intracellular membrane and microtubule behavior; centriole ultrastructure has been described previously (see, for example, Bettencourt-Dias et al., 2005; Rogers et al., 2008; Cunha-Ferreira et al., 2009).

*Drosophila* mitosis is semi-closed; prometaphase is not preceded by nuclear envelope breakdown (NEBD) but begins when nuclear envelope fenestrations at the cell poles allow penetration of astral MTs into the nucleus (Stafstrom and Staehelin, 1984; Debec and Marcaillou, 1997). Consistent with this notion, we found that approximately one-half of prometaphases were surrounded by nuclear membranes. These cells were real prometaphases, as they exhibited both compact chromatin patches and microtubules within the nucleoplasm. According to their morphology, we subdivided prometaphases into 4 classes: PM1 through PM4. To determine the temporal sequence of these classes we considered several parameters: the chromatin compaction, the MT arrangement, the structure of kinetochores and the intracellular membrane organization. In addition, we considered perinuclear lamin behavior, which is described below. Metaphases, were recognized based on the same parameters and, in many cases, on the examination of serial sections, and were included in a single class. Anaphases were distinguished into early and mid-late, and telophases were subdivided into early-mid and late.

### Prometaphase

We assigned 12 prometaphases to the PM1 class. To define the characteristic features of this class, we examined serial transverse sections through the chromosomes of one cell, and 11 longitudinal or slightly oblique sections with respect to the spindle axis; for 2 of these 11 cells we generated serial sections (an example of these serial sections is shown in Figure S2A). Cells in the PM1 class are characterized by the presence of a continuous membrane envelope (except at nuclear fenestrations) around the nucleus, mainly made by a typical double nuclear membrane (DNM) but also by patches of quadruple membrane (QNM) formed by two closely apposed double membranes connected by a layer of electron-dense material of unknown nature. In all cases, the QNM did not represent more than 18% of the total nuclear envelope length. In cells where fenestrations in the nuclear envelope were visible (in 6 out of 11 longitudinally cut cells), the QNM was always located in these regions and exhibited ribosomes on both the outer and inner surface of the membrane (Figures 1A, 2A and S2A). In the remaining 5 cells, patches of QNM were clearly present but could not be associated with nuclear fenestrations, as these membrane openings could not be seen due to the sectioning plane. In 10 out of 12 PM1 cells, we observed patches of endoplasmic reticulum (ER) membranes all over the cytoplasm; their average length per cell section was 11.7 ± 1.6 μm, while the length of the membranes forming the nuclear envelope, (DNM plus 2 × QNM) was 23.7 ± 2.8 μm (Figure 2B). Analysis of MTs in the transversally cut cell indicated that at this stage of mitosis MTs are rather sparse and are not organized in bundles as in later stages (Figure S3A). In addition, in 2 out of 11 longitudinally cut cells we were able to see immature kinetochores (Figure S4).

**Figure 1.**
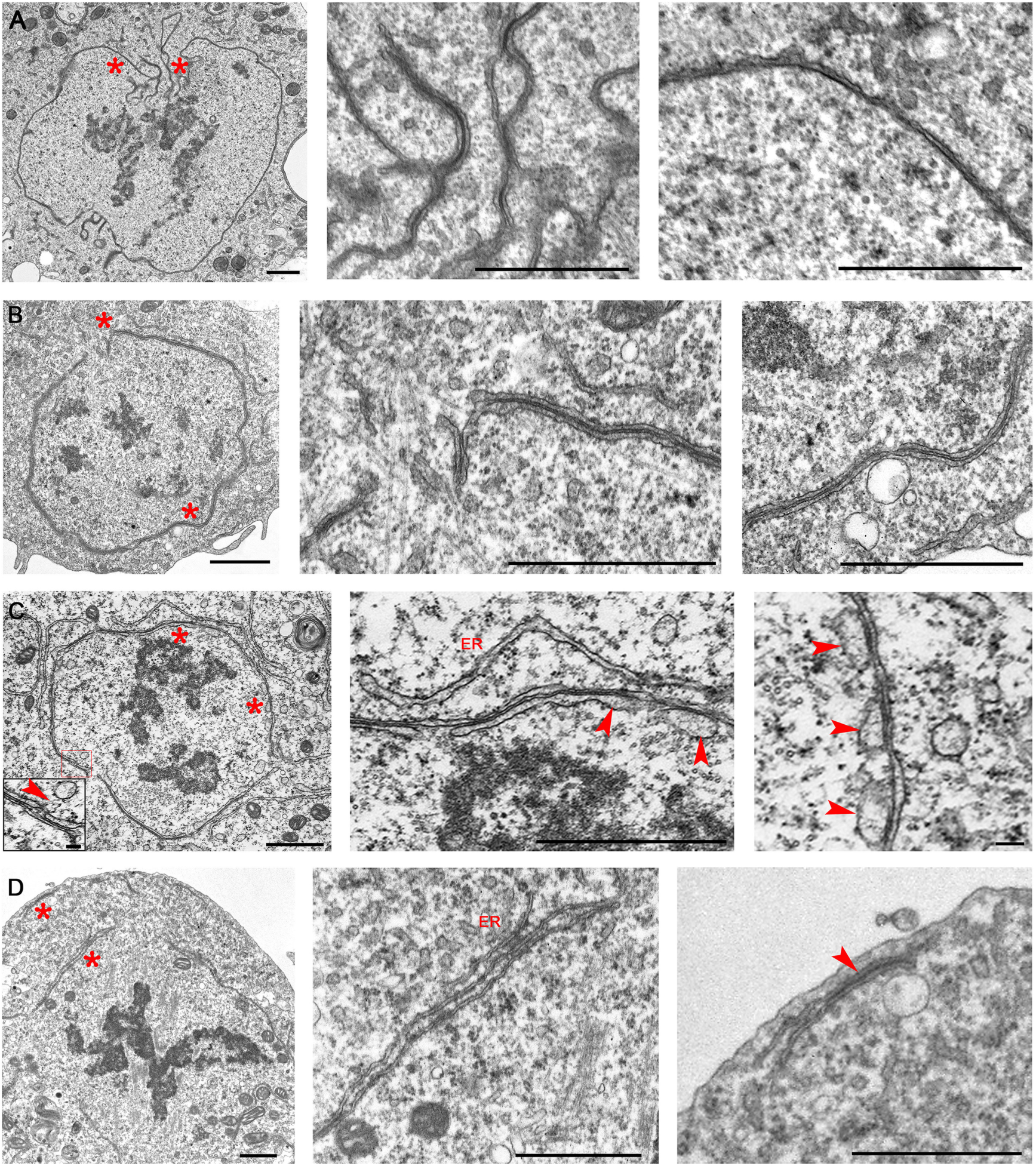
Prometaphase stages of S2 cells. (A) PM1 stage; note formation of QNM in the area of nuclear fenestration and the presence of a normal DNM along most nuclear envelope. (B) PM2 stage; note that almost all nuclear envelope is composed by QNM. (C) PM3 stage; note disassembly of the inner membrane of the QNM through vesiculation (inset; vesicles are indicated by arrowheads). (D) PM4 stage; note the almost complete disassembly of the QNM (remnants of the QNM are observed at the cell poles, arrowhead) and the appearance of ER membrane stacks. Asterisks indicate the cell regions shown at higher magnification in the central and rightmost images. Scale bars: 1 μm; except rightmost image of C, 0.1 μm.

**Figure 2.**
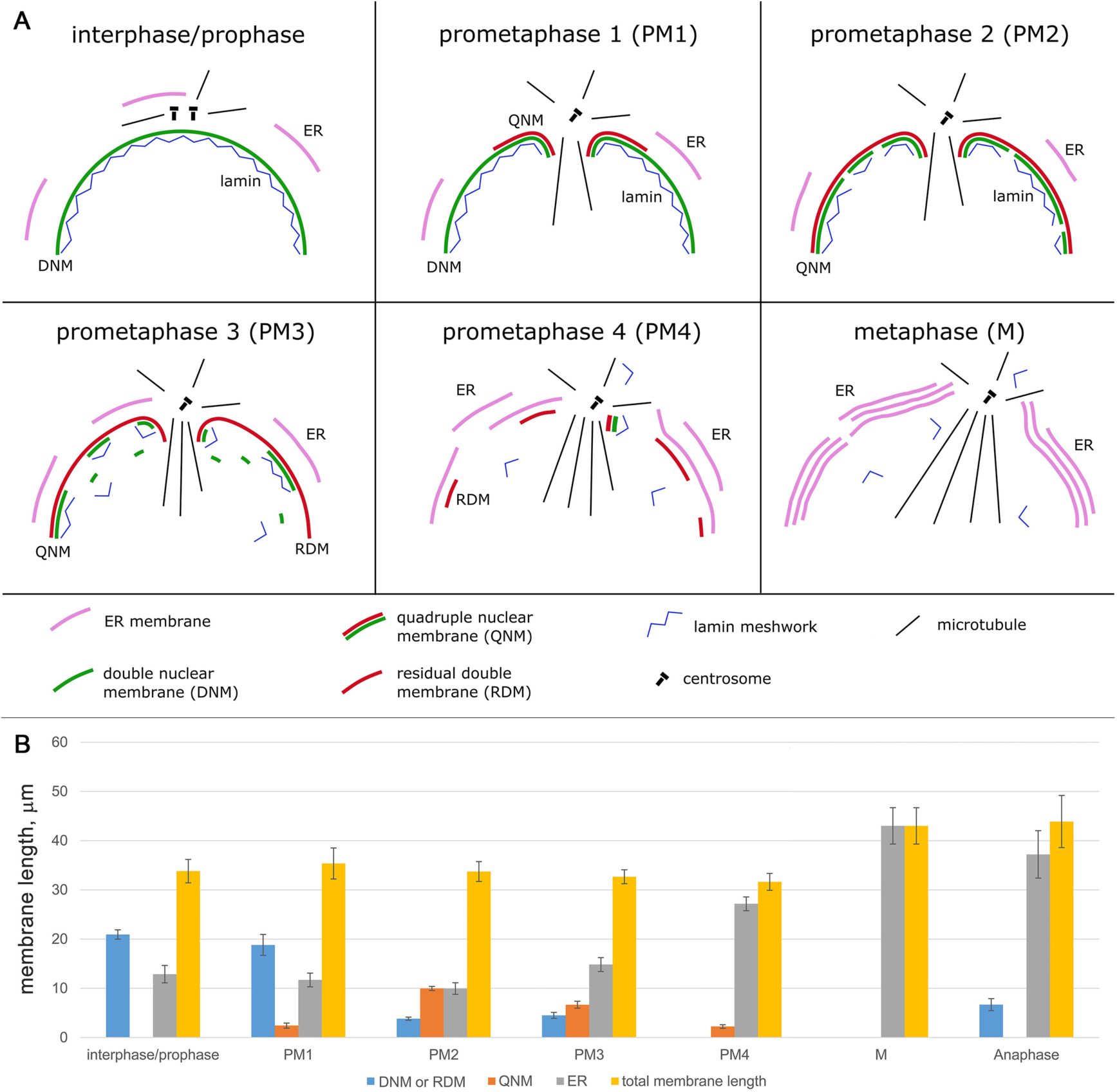
Schematic representation of intracellular membrane behavior during mitosis of S2 cells. (A) Membrane remodeling during prometaphase and metaphase. In PM1, a QNM begins to form in the proximity of the nuclear fenestration area, and during PM2 extends along the entire nuclear envelope. During late PM2 and PM3, the inner double membrane of the QNM becomes swollen and progressively disassembles through a vesiculation process, while a RDM persists around the nucleus. In PM4, only small patches of QNM are observed at the cell poles and the RDM disassembles. In metaphase cells, multilayer stacks of ER membranes form alongside the spindle.(B) Membrane length measured in 70 nm sections of interphase and mitotic cells. The DNM is present in interphase and prophase cells, in PM1 cells, and around some regions of anaphase chromosomes. In PM3 cells, the DNM is no longer present and the non-QNM portions of the nuclear envelope are made of RDM. To calculate the total membrane length we counted twice the QNM. Note that the total membrane length is similar from prophase through late prometaphase and increases in metaphase and anaphase cells. Error bars, SEM.

The prometaphases of the PM2 and PM3 classes show similar features and a continuum of morphological variations. They both exhibit a higher chromatin compaction compared to PM1 cells, and have a mostly continuous nuclear envelope that contains regions of QNM varying in length from 20% to 80% of the nuclear envelope. In some of the regions made of QNM, the inner membrane showed unevenly swollen traits and some discontinuities, suggesting that it is undergoing some form of decay (Figures 1B, 1C, 2A and S2B, C). In other nuclear envelope regions of variable length the inner membrane of the QNM had completely disappeared (Figures 1B, 1C, 2A and S2B, C), leaving a residual double membrane (henceforth RDM). We observed ribosomes associated with both the outer and inner surface of the QNM as well as with both surfaces of the RDM (Figures 1B and S2B, C).

We arbitrarily assigned to the PM2 class 17 cells based on the fact that their nuclear envelopes are formed by at least 60% of QNM. This class includes 1 cell represented by serial transversal sections through the chromosomes, and 16 longitudinally sectioned cells, 3 of which are represented by serial sections. In 13 out of these 17 PM2 cells, we observed patches of ER membranes; their average length per cell section was 10.0 ± 1.2 μm, while the length of the membranes forming the nuclear envelope, (2 × QNM plus RDM) was 23.8 ± 1.3 μm (Figure 2B). An analysis of the transverse section indicated that also in this stage of mitosis MTs are rather sparse, although some loose MT bundles were apparent (Figure S3B). In 4 out of the 15 cells examined we were able to see immature kinetochores (Figure S4).

We assigned to PM3 stage 18 cells with nuclear envelope containing less than 60% of QNM. This class includes 2 cells cut transversally through the chromosomes, one of which was serially sectioned; and 16 longitudinally cut cells, 9 of which are represented by serial sections. In these PM3 cells, the inner membrane of the QNM was usually more swollen (80 ± 5 nm against 40 ± 1 nm, post hoc comparison using Tukey HSD test: p < 0.01) and discontinuous that in PM2 cells, suggesting that it was undergoing a degradation process through vesiculation (Figures 1C, 2A and S2C). 16 out of the 18 cells in the PM3 class showed layers of ER membranes surrounding more or less extensive regions of the nuclear envelope (formed by QNM and RDM). In these 16 cells, the average length of the ER membranes per cell section was 14.8 ± 1.4 μm, while the length of the membranes forming the nuclear envelope, (2 × QNM plus RDM) was 17.8 ± 1.8 μm (Figure 2B). When the ER membrane layers were sufficiently long, their curvatures tended to conform to that of the nuclear envelope (Figures S2B and S3C). However, in contrast with the typical QNM appearance, ER membranes rarely showed close apposition to either the QNM or the RDM. We observed only a few cells where this type of adhesion appeared to occur in a limited portion of the nuclear envelope (Figure S5). An analysis of MTs in transverse sections indicated that in this stage of mitosis there is a diminution of isolated MTs compared to the PM2 class (Figure S3C). Some MT bundles were visible, but these bundles were fewer compared with the bundles observed in the following stage of prometaphase (PM4) and in metaphase cells. In 11 out of the 18 cells examined we were able to see immature kinetochores (Figure S4).

We included 24 cells in the PM4 class, which includes transverse sections from 3 cells (1 serial), and longitudinal sections from 21 cells (6 serial). These cells are characterized by compact chromatin and the disappearance of almost all QNM and RDM. However, most spindles of the PM4 cells (19 out of 24) were partly enveloped by one or more irregular layers of ER membranes comprising from 1 to 3 layers running alongside with little or no overlapping (Figures 1D). In the 19 cells with ER membranes, the average length of these membranes was 27.2 ± 1.4 μm per cell section, while the length of the remaining 2× QNM plus RDM was 5.8 ± 1.6 μm (Figure 2B). An analysis of MTs in transverse sections indicated that in PM4 cells most MTs are organized in bundles containing from 4 to 45 MTs (Figure S3D). In addition, in a substantial fraction of longitudinal sections (8 out of 16) we were able to see mature kinetochores, most of them showing end-on MT attachment (Figure S4).

### Metaphase

The 37 cells assigned to this class include 7 cells cut transversally through the chromosomes (4 which are represented by serial sections), and by 30 longitudinally cut cells (11 of which were serially sectioned). These cells are characterized by the alignment of highly compacted chromosomes at the cell equator and by the presence of robust MT bundles often ending at the kinetochores (Figures 3 and S6). In 26 of the 30 longitudinal sections examined, the spindles and chromosomes were surrounded by one or more discontinuous layers of ER membranes located near the cell poles, at the cell equator or at both the poles and the equator. Most of these metaphases showed 2-3 parallel ER membrane layers but some of them displayed from 4 to 7 membrane layers (Figures 2, 3 and S6A; the cell shown in Figure 3 exhibits 7 parallel layers of ER membranes). In sections showing multiple ER membrane layers, these membranes run alongside the spindle with little or no overlapping, and never formed a quadruple membrane comparable to that observed in prometaphase cells. Examination of multiple sections of the same cell showed that the ER membrane layers that surround the metaphase spindles are organized in discontinuous and irregular stacks, which contain a variable number of membranes (see Figure S2D as an example). Thus, the metaphase spindle is not isolated by a “spindle envelope” but has many regions that are directly in contact with the mitotic cytoplasm. In the 26 cells showing ER membranes, the average membrane length per section was 43.0 ± 3.7 μm (Figure 2B).

**Figure 3.**
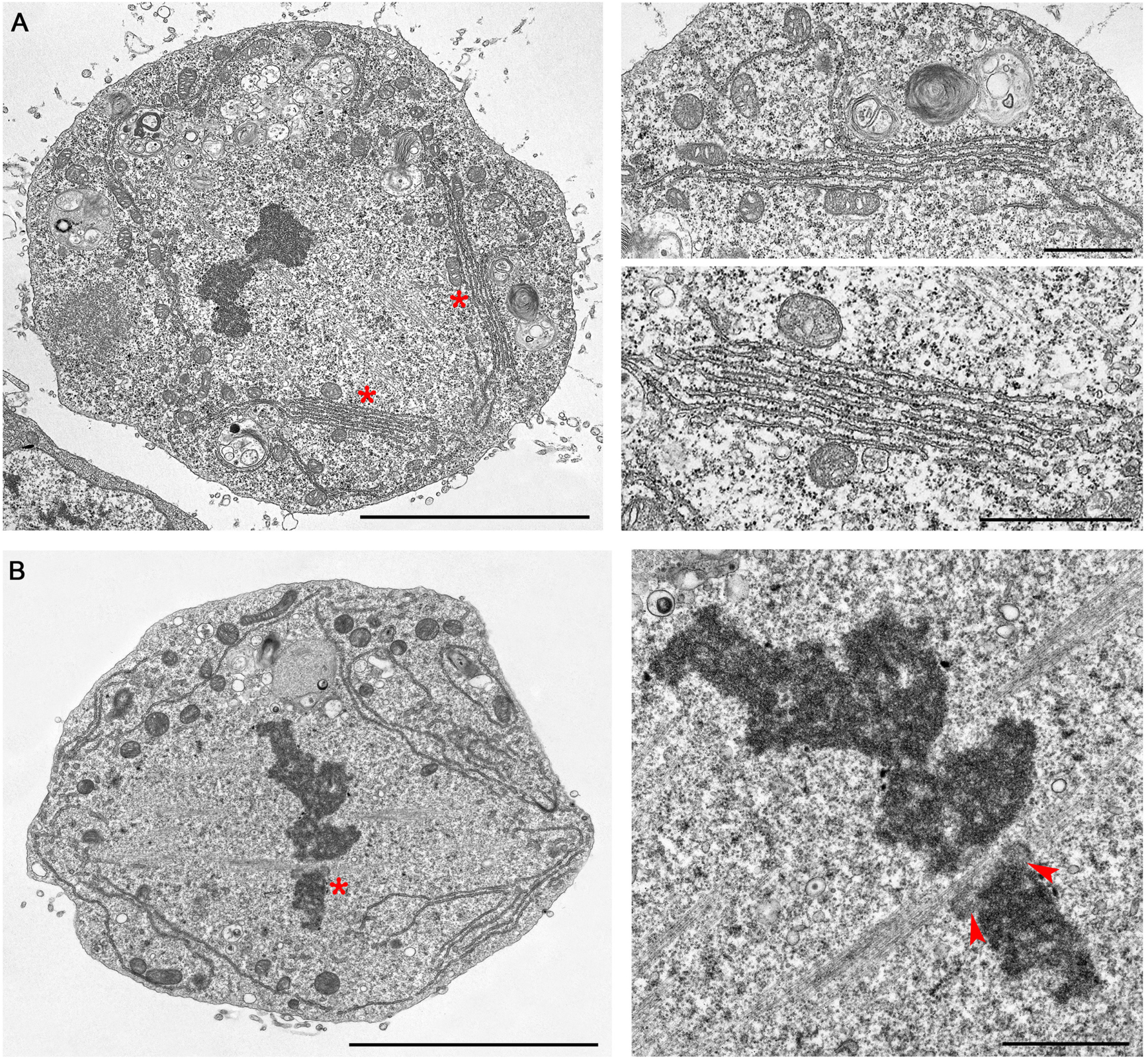
Examples of S2 cell metaphases. Note the stacks of parallel ER membranes in A, and the kinetochores with k-fibers (arrowheads) in B. Asterisks indicate the cell regions shown at higher magnification in the images on the right. Scale bars: Left images 5 μm; right images 1 μm.

Examination of the transverse sections showed that metaphase MTs are mainly organized in bundles; some of these bundles end directly on the kinetochores while others appear to run alongside the kinetochores (Figure S6).

### Anaphase

We examined only longitudinal or slightly oblique sections of anaphases. We sectioned 12 early anaphases and 9 mid-late anaphases B cells; we made serial sections of 7 and 5 of these cells, respectively. We classified as early anaphases cells in which the sister chromatids are separated and have started moving towards the poles, although the cell has still an oval shape comparable to that of metaphase cells (Figure 4A and S7). The MTs of early anaphase cells were still arranged in bundles; in 6 out of the 11 cells examined we observed clear kinetochore structures (Figure 4A).

**Figure 4.**
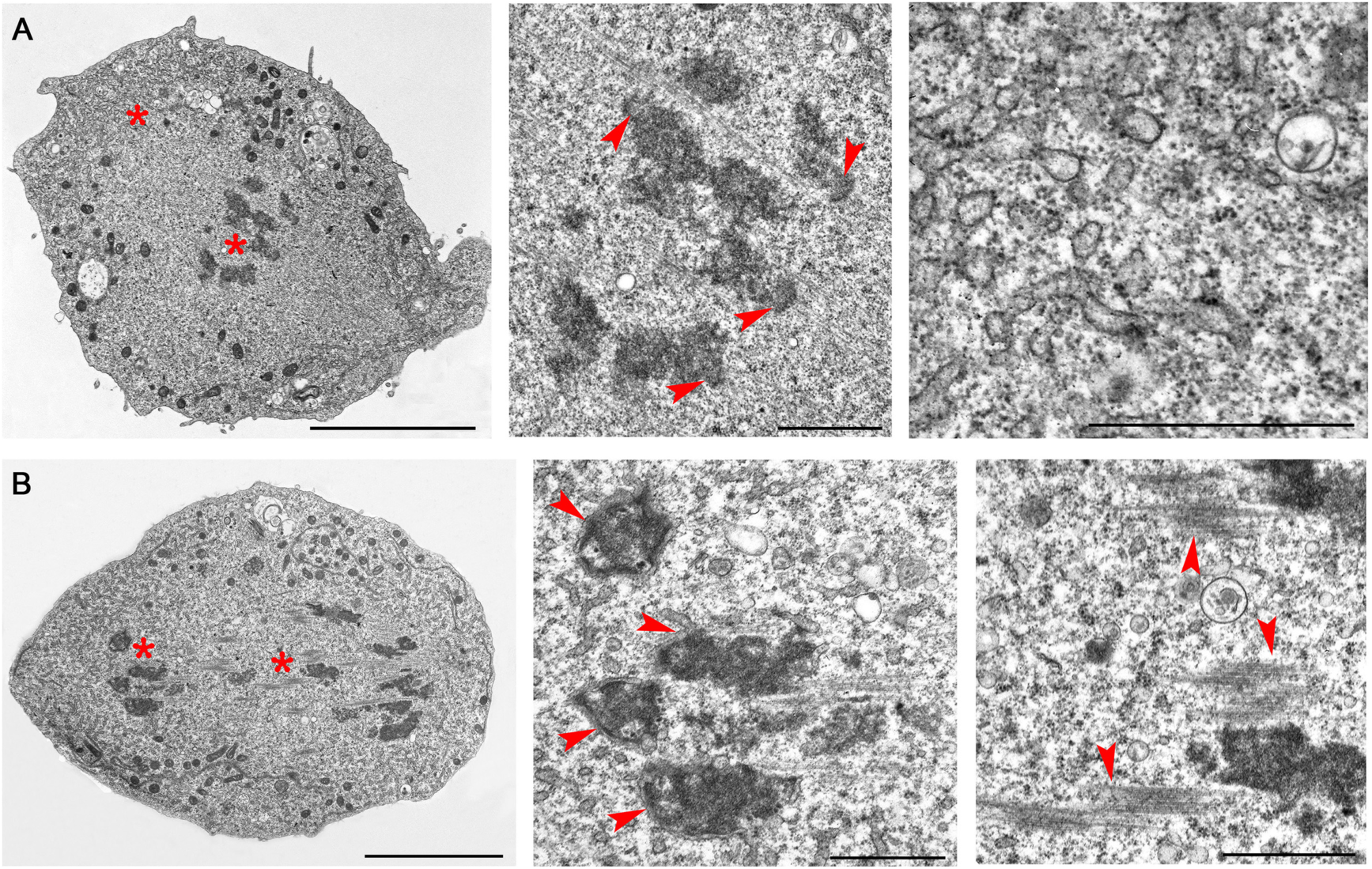
Examples of S2 cell anaphases. (A) Early anaphase cell showing initial separation of the sister chromatids. Note the kinetochores of the separating sister chromatids (arrowheads) and the vesicles at the spindle poles. (B) Mid-late anaphase. Note the formation of a double membrane around the chromosomes (arrowheads) and the MT bundles in the center of the cell. Asterisks indicate the cell regions shown at higher magnification in the central and rightmost images. Scale bars: leftmost image 5 μm, central and rightmost images 1 μm.

In the mid-late anaphase cells, the two sets of separating chromatids have reached the spindle poles or are located very close to them, and the cells exhibit an elongated shape compared to metaphase cells (Figure 4B and S7). In 8 out of the 9 mid-late anaphase cells, we observed large but discontinuous patches of double membrane surrounding the chromosomes; only 4 out of the 11 early anaphase cells examined showed very small patches of chromosome-associated membranes (Figures 4B and S7). Kinetochores and MT bundles connecting the chromosomes with the spindle poles were never observed in mid-late anaphase cells. However, longitudinal sections of these cells showed bundles of MTs traversing the sets of migrating chromosomes and/or placed between these sets in the center of the cell (Figures 4B and S7). The latter MT bundles will probably give rise to the telophase central spindle.

In both early and mid-late anaphase cells, the ER membrane were mostly confined in the polar regions of the cell but also formed single layers alongside the central part of the spindle; these ER membranes were shorter and often undulated compared to those seen in metaphase cells (Figure 4 and S7). We measured the ER membranes also in sections of anaphase cells (early and mid-late) and found that their average length was 37.2 ± 4.8 μm per section (Figure 2B). In mid-late anaphase cells, the average length of DNM forming around the chromosomes was 6.7 ± 1.2 μm (Figure 2B). The polar regions of anaphase cells also displayed many membrane vesicles (Figures 4 and S7) that are likely to be intermediates in the Golgi reassembly process. In S2 cells, like in mammalian cells, the Golgi apparatus disassembles upon entry of the cells into mitosis, giving rise to large number of fragments that disperse into the cytoplasm; these fragments progressively reaggregate during anaphase and telophase to eventually reassemble the Golgi stacks in the daughter cells (Stanley et al., 1997; Kondylis et al., 2007; Shima et al., 1997).

### Telophase

As for anaphases, we examined only longitudinal or slightly oblique sections containing both presumptive daughter cells. We sectioned 7 early-mid telophases and 8 late telophases; we made 3 serial sections for each of these stages. Early-mid telophase figures were characterized by the decondensation of the chromosomes, which are almost completely surrounded by a DNM. These cells, in addition to an initial cleavage furrow, showed deformations of the surface with frequent bulges protruding from both the equatorial region and the cell poles (Figures 5A and S8B). These cell protrusions were not observed in prometaphase, metaphase and anaphase cells. Early-mid telophases also exhibited separate MT bundles between the cell poles, in the region where the central spindle is forming (Figure S8A).

**Figure 5.**
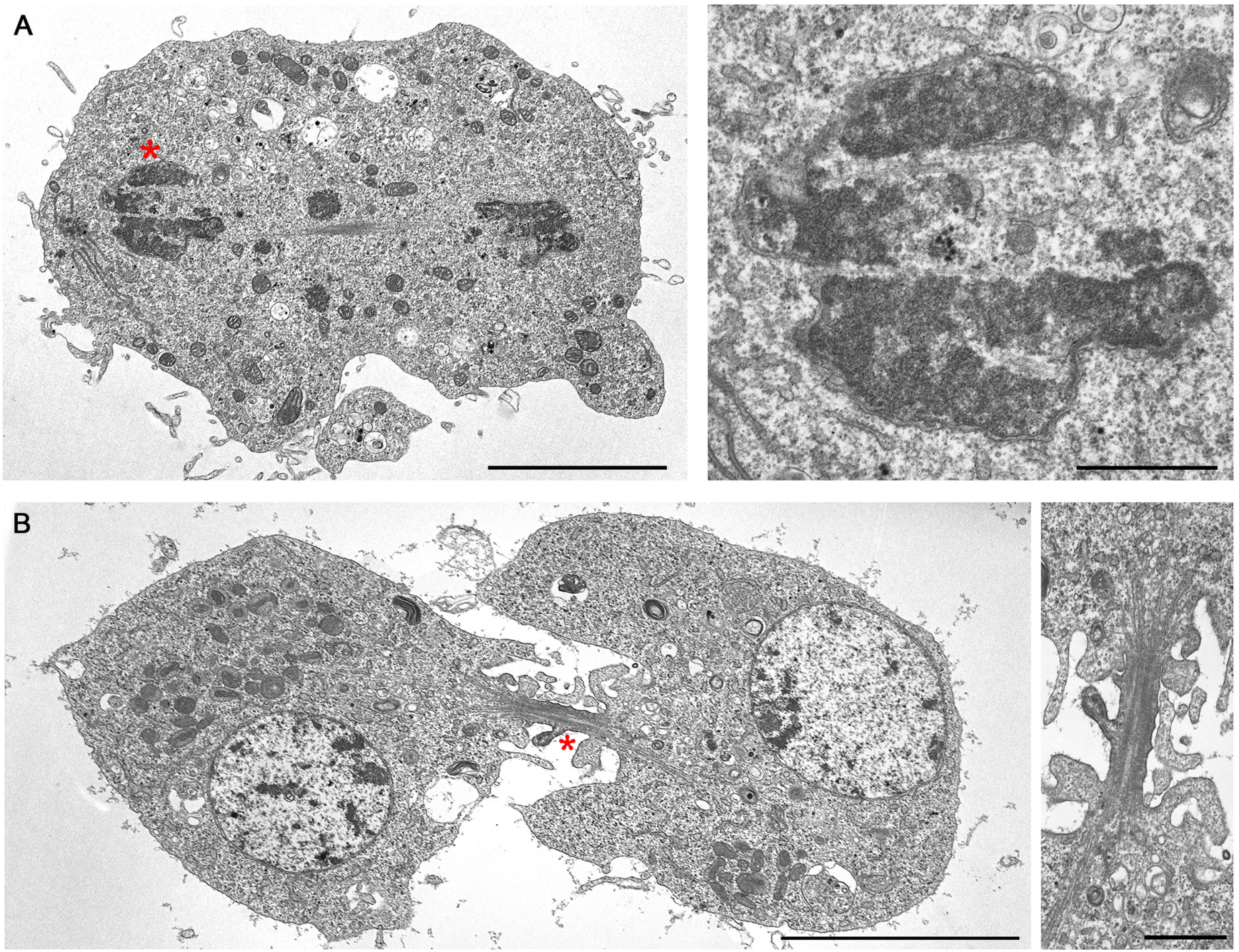
Examples of S2 cell telophases. (A) Early telophase with the chromosomes almost completely surrounded by a double membrane. Note the bulges protruding from both the equatorial region and the cell pole. (B) Late telophase with fully reformed nuclei. Note the multiple protrusions emanating from the intercellular bridge. Asterisks indicate the cell regions shown at higher magnifications. Scale bars: A, left image 5 μm, right image 1 μm; B, left image 5 μm, right image 1 μm.

Late telophases are characterized by fully reformed nuclei containing uncondensed chromatin surrounded by a continuous DNM. The two presumptive daughter cells were connected through an intercellular bridge that contained bundled MTs. In some cells, the plus ends of the MT bundles coming from the opposite daughter cells were clearly overlapping at the center of the intercellular bridge (Figure S8C, D). One interesting feature of the late telophase cells was the presence of many protrusions emanating from the cell surface. Some of these protuberances were rather large but others were small and often exhibited a tube-like appearance; the small protrusions were mostly emanating from the intercellular bridge (Figures 5B and S8C, D). Large membrane bulges have been also observed with a light microscope in telophase of normal S2 cells fixed with formaldehyde/PBS and stained for actin, tubulin and DNA, and their frequency and size were greatly enhanced in anillin-depleted cells (Somma et al., 2002). It is likely that both the pole- and the intercellular bridge-associated protrusions are dependent on changes in the cell cortex contractility and the extensive membrane remodeling that occurs during cytokinesis (Sedzinski et al, 2011; Cabernard, 2012; Schiel and Prekeris, 2013).

### Membrane behavior during S2 cell mitosis

Extensive literature supports the notion that the double membrane that forms the nuclear envelope in interphase cells is continuous to the endoplasmic reticulum (ER). The outer nuclear membrane is connected to the ER through tubules or sheets and shares with the ER many proteins and lipids. In contrast, the inner nuclear membrane is biochemically distinct from the outer and contains hundreds of unique components, many of which are thought to interact with the nuclear lamina and the chromatin (Schirmer and Gerace, 2005; Crisp and Burke, 2008; Güttinger et al., 2009; Korfali et al., 2012). We found that interphase and prophase cells exhibit a DNM that surrounds the nucleus and a few patches of ER membranes that do not appear to be closely associated to the DNM. However, starting from early prometaphase, the membrane systems of S2 cells exhibit a series of dramatic transformations, which are schematically summarized in Figure 2A.

Our results clearly show that at the beginning of prometaphase the nuclear envelope exhibits small regions made of quadruple membrane (QNM); the nuclear fenestration areas were always characterized by the presence of QNM. Following this initial appearance, the QNM forms along the entire nuclear envelope. Our data do not allow us to envisage a precise mechanism for QNM formation. Clearly, QNM assembly requires the formation of an additional double membrane along the DNM. In principle, this additional membrane could be synthesized *de novo* either at the inside or the outside of the DNM. Alternatively, it could be a pre-existing ER membrane, which would first become closely apposed and then attach to the DNM. We favor this second possibility for several reasons. First, as mentioned earlier, there is continuity between the nuclear membrane and the ER, which are connected by tubules or sheets, a condition that would facilitate conjoining of the two double membranes. Second, we observed several early prometaphase cells showing ER membrane running alongside to the DNM, and a few examples of close apposition (Figures S2B, C and S5) of the ER membranes to the DNM, two findings in line with the second alternative. Third, there are several examples of apposition of ER membranes reported in the literature. The most pertinent one concerns *Drosophila* syncytial embryos, where during prometaphase a second ER membrane forms around the double nuclear membrane (Stafstrom and Staehelin, 1984). In another example, downregulation of the lipin homologue of *C. elegans* led to the formation of an extra ER layer in close association with the nuclear membrane and interfered with nuclear envelope disassembly (Bahmanyar et al., 2014). Stacked layers of ER membranes have been also observed in vertebrate cells overexpressing certain ER membrane proteins (Wright et al., 1988; Snapp et al., 2003) or infected with the African swine fever virus (Windsor et al., 2012). However, in all these systems the ER sheets were not attached and did not show the type of adhesion observed in the QNM of S2 cells. Patches of QNM similar to those seen in S2 cells have been observed only in mammalian tumor cells undergoing mitosis but not in normal dividing cells (see for example Chang and Gibley, 1968; Parmley et al., 1976 and references therein). It has been also suggested that the QNM observed in these tumor cells is generated by the attachment of ER segments to the nuclear envelope (Parmley et al., 1976).

We have shown that at the fenestration sites the nuclear membrane exhibits multiple folds and invaginations that are mostly comprised of QNM. This dramatic deformation of the nuclear membrane is likely to be caused by the astral MTs that come into contact with the nucleus. Deep microtubule-dependent invaginations in the nuclear envelope have been also observed in vertebrate cells before nuclear envelope breakdown (NEBD), and it has been suggested that MT-nuclear envelope interactions trigger NEBD. However, NEBD can also occur in the absence of MTs suggesting the existence of a MT-independent pathway for NEBD (reviewed by Güttinger et al., 2009; Fernández-Álvarez and Cooper, 2017). These findings prompted us to investigate whether QNM formation requires an interaction with the astral MTs. To answer this question we performed RNAi against the *cnn* gene that encodes a centrosome component essential for MT nucleation (Li et al., 1998). Consistent with previous studies (Somma et al., 2008) we found that RNAi-mediated Cnn depletion completely inhibits aster formation in S2 cells. TEM analysis of Cnn-depleted cells revealed that several prometaphase cells exhibit a QNM (Figure S9), indicating that formation of this peculiar membrane structure does not depend on interactions between the DNM and the astral MTs.

With progression through prometaphase, the inner double membrane of the QNM becomes swollen and progressively disassembles through the apparent formation of cisternae that detach from the outer membrane of the QNM. Interestingly, nuclear envelope-derived cisternae have been also observed in mammalian cells undergoing NEBD (Waterman-Storer et al., 1993; Salina et al., 2002; Cotter et al., 2007). The finding that both the inner component of the QNM and the mammalian nuclear envelope disassemble through vesicle formation supports the hypothesis that the inner membrane of the QNM corresponds to the DNM of interphase and PM1 cells. Disassembly of the inner membrane of the QNM is followed by the appearance of several openings in the nuclear envelope and the progressive transition from a closed to an open form of mitosis.

The measure of the length of the different membrane types (ER membranes, DNM, QNM, and RDM) in 70 nm sections showed that total membrane length (counting twice the length of the QNM) in interphase, PM1, PM2, PM3 and PM4 cells is comparable. In contrast, total membrane length in metaphases and anaphases is approximately 1.5-fold greater than in interphase or prometaphase cells (Post hoc comparisons using the Tukey HSD test, p < 0.05) (Figure 2B). This finding suggests that during prophase and prometaphase there is an extensive membrane remodeling with respect to interphase, but no biosynthesis of new membrane. In contrast, during the short period that elapses between late prometaphase and metaphase (~ 5 minutes; based on our movies of S2 cell division; see Renda et al., 2017), additional membrane is synthesized, a process that is probably related with the need of membrane for the generation of the daughter cells during cytokinesis.

An analysis of nuclear pore complex (NPC) distribution in 70 nm sections showed that the interphase nuclei contains 21.8 ± 1.4 NPCs, with an average of one NPC per 0.98 μm of DNM. In the PM1 stage, the average number on NPCs in the DNM was drastically reduced (4.7 ± 0.9), with an average of one NPC per 4.4 μm of DNM. The QNM was consistently devoid of NPCs crossing both double membranes. However, NPC-like structures were rarely observed in the external double membrane of the QNM (Figure S10A) and the RDM of PM3 and PM4 cells, and even in the ER membranes approaching the nuclear envelope (Figure S10B). NPCs have been previously observed in ER membranes that form the annulate lamellae in *Drosophila* syncytial embryos (Stafstrom and Staehelin, 1984). The ER membrane layers alongside metaphase and anaphase spindles of S2 cells did not exhibit any NPC-like structures. NPCs reassembled concomitant with membrane formation around late anaphase B and telophase nuclei (Figure S10C, D). Thus, as occurs in mammalian cells (Güttinger et al., 2009; Fernández-Álvarez and Cooper, 2017), and consistent with previous work on *Drosophila* syncytial embryos (Kiseleva et al., 2001), NPCs of S2 cells disassemble at the onset of mitosis and reassemble during telophase.

### Lamin behavior and the role of the QNM during S2 cell mitosis

To integrate TEM observations, we stained with anti-lamin and anti-GFP antibodies dividing S2 cells that express tubulin-GFP, and examined them by confocal microscopy (Figure 6). Remarkably, the analysis of optical sections of these cells permitted visualization of the different stages of prometaphase identified by TEM analysis. We analyzed 160 prometaphase cells; 16% displayed a continuous lamin envelope except at fenestrations penetrated by MTs (Figure 6B). These cells most likely correspond to the PM1 and PM2 stages. 37% of the prometaphases showed an incomplete lamin envelope (Figure 6C, D) and are likely to correspond to the PM3 stage. The remaining 49% of the prometaphases examined displayed some remnants of the lamin envelope and small patches of lamin dispersed throughout the cytoplasm (Figure 6E, F); these cells are most likely in the PM4 stage. In metaphase cells, chromosomes were not surrounded by any clear lamin-enriched structure, suggesting that in S2 cells the lamin envelope disassembles in late prometaphase. Thus, it appears that lamin envelope disintegration occurs in the PM4 stage at the transition between prometaphase and metaphase.

**Figure 6.**
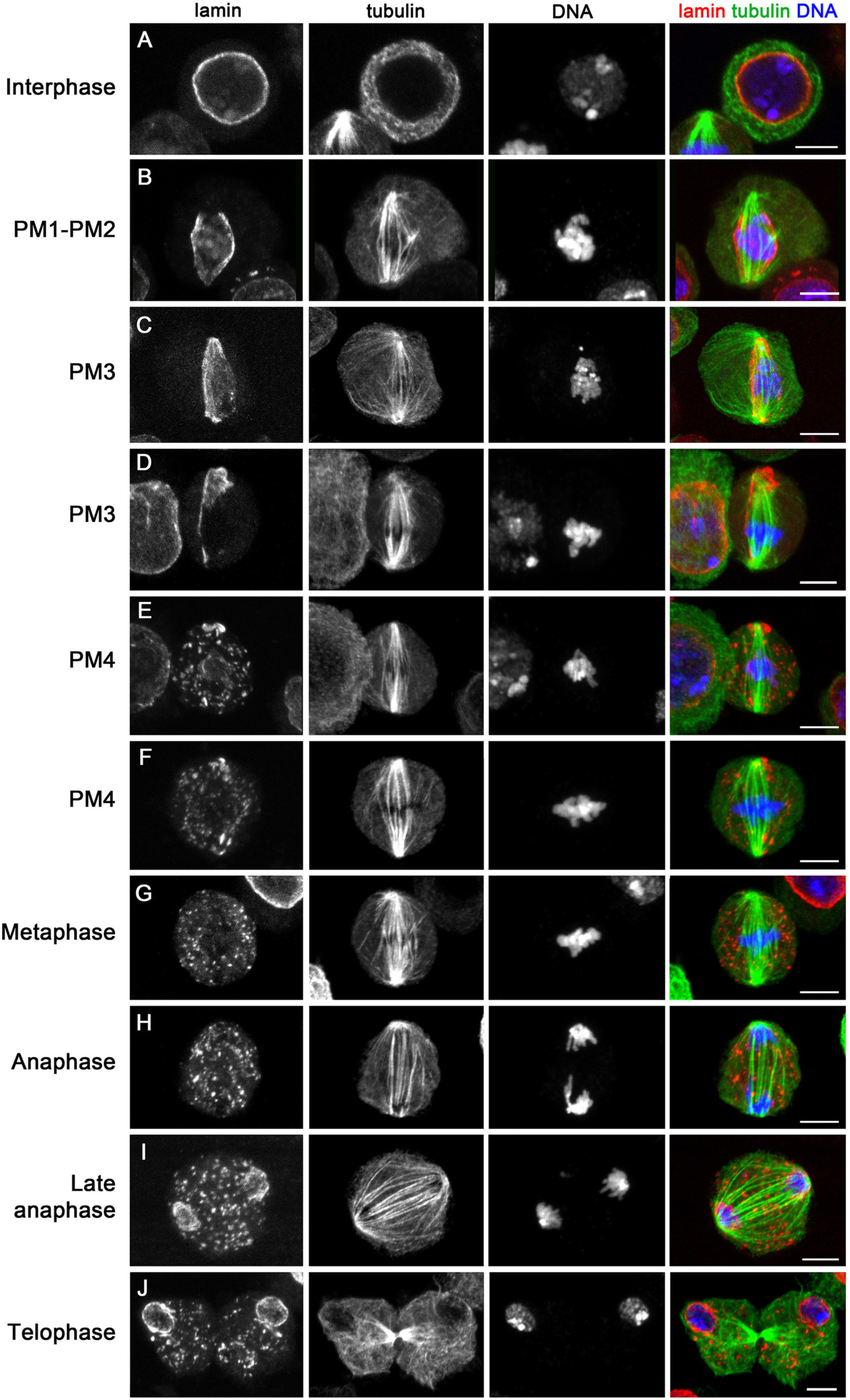
Lamin immunostaining recapitulates TEM observations. Cells were stained for lamin, tubulin and DNA. A lamin meshwork encases the nucleus in interphase and early prometaphase cells (PM1 and PM2), begins to disintegrate in mid-prometaphase (PM3), and it is completely disassembled in late prometaphase (PM4) and metaphase cells. Note that fragments of the nuclear lamina persist from prometaphase till late telophase. Scale bars: 5 μm.

Interestingly, after prometaphase, lamin-enriched speckles persisted through metaphase, anaphase and telophase (Figure 6 G-J). These lamin speckles have been observed previously in S2 cells (Schweizer et al., 2015) but not in *Drosophila* embryonic or brains mitoses and in male meiosis (Harel et al., 1989; Paddy et al., 1996; Katsani et al, 2008; Fabbretti et al., 2016). We do not know the composition and the function of these speckles, which were not detected by TEM analysis. They might be aggregations of lamin, other nuclear envelope proteins and membrane that did not completely disassemble to facilitate reformation of the nuclear envelope at telophase. Indeed, starting from late anaphase and throughout telophase, we observed a lamin envelope surrounding the two daughter nuclei (Figure 6I, J). However, the lamin speckles persisted also after initial reformation of this nuclear envelope raising the possibility that they could be used for nuclear envelope expansion in late telophase or during the ensuing G1.

Collectively, our results suggest the hypothesis that *Drosophila* S2 cells evolved a QNM to maintain a closed mitosis up to mid-late prometaphase, to facilitate MT-kinetochore interactions during early prometaphase. Our observations suggest that the inner double membrane component of the QNM corresponds to the vertebrate nuclear membrane, which disintegrates at the beginning of prometaphase concomitantly with its associated lamin meshwork (NEBD). The formation of a QNM would maintain the continuity of the nuclear envelope even when the inner double membrane component of the QNM disassembles, thus postponing NEBD from the beginning of prometaphase to mid-late prometaphase (stage PM4). The lamin behavior during prometaphase (Figure 6) raises the question of whether the stability of the perinuclear lamin meshwork requires an interaction with the inner component of the QNM or can also be stabilized by its outer component (the RDM). Our light microcopy observations (Figure 6) appear to support the first alternative. However, further studies will be required to define the properties of the two double membranes of the QNM.

### Relationships between mitochondria and ER membranes

Studies carried out in both *Drosophila* and human cells have shown that membranous organelles (e.g., mitochondria) are excluded from the spindle region through a microtubule independent mechanism (McIntosh and Landis, 1971; Schweizer et al., 2015, and references therein). It has also been shown that soluble tubulin and proteins such as Megator and Mad-2 accumulate in the nucleoplasm soon after nuclear fenestration at the beginning of prometaphase, and remain within the nucleus for approximately 10 minutes before spreading into the cytoplasm (Schweizer et al., 2015). To explain this finding, it has been suggested that during early mitosis these proteins are confined into the nucleoplasm by “spindle envelope” membranes that still surround the nucleus.

Our analyses showed the presence of a complex membranous system encasing the chromosomes during the early stages of prometaphase, and that this system becomes discontinuous after the PM3 stage. Nevertheless, our TEM photographs revealed that mitochondria are mostly excluded from the spindle/nuclear area not only in the PM1, PM2 and PM3 prometaphase stages but also in during the PM4 stage and the subsequent mitotic stages. We found that mitochondria are mostly associated with ER membranes that discontinuously surround the spindle during late prometaphase (PM4), metaphase, anaphase and early telophase. These organelles were often found in close proximity or attached to the ER membranes on both their inner and outer sides with respect to spindle (Figures 3, 4, S2D, S6, S7, S8B); sometimes they were anchored to these membrane like bulbs on a garland (Figure S8B). An intimate interconnection between mitochondria and ER has been observed in yeast and shown to be crucial for proper mitochondria distribution during mitosis. However, it is unclear whether an ER and mitochondria also associate during animal cell mitosis (reviewed in Rowland and Voeltz, 2012). Our results strongly suggest that the mitochondria and the ER physically interact during S2 cell mitosis, and confirm the exclusion of these organelles from the spindle area (Schweizer et al., 2015). However, our observations indicate that ER membranes do not form a barrier against the invasion of the spindle region by mitochondria as suggested previously (Schweizer et al., 2015), but instead constitute a trap that captures these organelles, preventing their free diffusion within the dividing cells.

### Membrane behavior during mitosis in different *Drosophila* tissues

The most studied *Drosophila* mitotic system is the syncytial embryo, characterized by a series of extremely rapid and synchronous mitotic divisions. Membrane behavior in this system is similar but not identical to the one just described for S2 cells. Also in embryonic divisions a second membrane is added just outside the nuclear membrane; this process begins in prometaphase near the fenestrated areas at the spindle poles and is completed at metaphase with the formation of two membrane layers that surround the entire spindle with the exception of the poles. These double membrane layers, often designated as spindle envelope, persist during metaphase and anaphase, contain ER markers, exhibit numerous irregular fenestrae but are never as closely apposed as the two double membranes that comprise the S2 cell QNM (Stafstrom and Staehelin, 1984; Bobinnec et al., 2003; Frescas et al., 2006). The regions of single nuclear membrane around the embryonic prometaphase nuclei exhibit disassembling NPCs but the two membrane layers of metaphase nuclei are virtually devoid of NPCs, which reassemble only in the membrane that surrounds telophase nuclei (Stafstrom and Staehelin, 1984; Kiseleva et al., 2001). Finally, in contrast with S2 cells where the nuclear lamina disintegrates during prometaphase, in embryos the lamina surrounds spindles till late metaphase and dissolves just before anaphase (Paddy et al., 1996; Katsani et al., 2008). Thus, it appears that embryonic mitosis is more “closed” than S2 cells mitosis, consistent with the fact that embryonic nuclei are not surrounded by a plasma membrane.

A spindle envelope similar to that seen in embryonic mitoses has been also observed in dividing neuroblasts of *Drosophila* larval brains; like in embryos this membranous structure persists till late anaphase (TEM observations by A.T.C. Carpenter, personal communication). In addition, it has been shown that from metaphase to late anaphase of brain neuroblasts the phosphatidylinositol transfer protein (PITP) Giotto is enriched at an intracellular structure that for location and shape corresponds to the spindle envelope (Giansanti et al., 2006). PITPs are enriched at cell membranes where they bind lipid monomers, facilitating their transfer between different membrane compartments (Allen-Baume et al., 2002). Consistent with these results, *in vivo* studies have shown that GFP-lamin marks the nuclear envelope from prophase till anaphase, although the anaphase GFP signal is weaker than those observed in prometaphase and metaphase cells (Katsani et al., 2008). These observations suggest that neuroblast mitosis has a similar membrane behavior as mitosis in syncytial embryos. Thus, S2 cell mitosis appears to be more “open” than both neuroblast and syncytial embryo mitosis.

The few studies performed in S2 cells have not provided details on the intracellular membrane behavior during mitosis; it has been only reported that a membrane-based nuclear envelope disappears just prior to metaphase plate formation (Maiato et al., 2006). A study on the heat shock effects in Kc cells, another *Drosophila* cell line of embryonic origin, showed that the spindles of cells not treated with heat are surrounded by two membrane layers that run adjacent to each other but do not exhibit areas of close apposition (Debec and Marcaillou, 1997). Thus, they are somewhat similar to the paired double membranes that encase embryonic divisions. “Spindle envelopes” containing a variable number of membrane layers have been described in *Drosophila* spermatocytes (Tates, 1971; Fuller, 1993) and in mitotic and meiotic spindles of several Diptera and Lepidoptera (Wolf, 1990; Wolf, 1995). However, the complex intracellular membrane behavior we observed in S2 cell mitosis has not been described in any of these systems. Moreover, published images did not show closely apposed membrane layers comparable to QNM we observed in S2 cells (Tates, 1971; Wolf, 1990; Wolf, 1995, Debec and Marcaillou, 1997).

### MT behavior and MT-kinetochore interactions in S2 cells

To analyze MT behavior and MT-kinetochore relationships we examined both single and serial sections of mitotic cells cut either alongside or transversely with respect to the spindle axis. We first considered transverse sections through prometaphase and metaphase chromosomes. The PM1, PM2, PM3, PM4 and metaphase stages were recognized for the degree of chromatin compaction and for the intracellular membrane pattern. In early stages of prometaphase, MTs are rather sparse but tend to aggregate into bundles as prometaphase proceeds (see Figures S3 for overall examples of MT distribution in transverse sections; detailed examples are shown in Figures 7A and 8B). To characterize the MT bundles at different mitotic stages, we measured the average distance between the MT centers within each bundle (see Material and Methods for the measuring procedures). We considered as MT bundles geographically isolated MT aggregates in which the average distance between the MTs is between 35 and 50 nm; aggregates of 2 or 3 MTs were not considered as bundles. This analysis showed that the distance between bundled MTs decreases from early prometaphase to metaphase (Figure 7B). The average inter-MT distances observed in the PM1, PM2 and PM3 bundles are significantly different (post hoc comparisons using the Tukey HSD test, p < 0.01), whereas the inter-MT distances within the PM3, PM4 and metaphase bundles are not significantly different. In addition, with progression from early prometaphase to metaphase there is an increase in the frequencies of large bundles containing 10-15, 16-21, and >21 MTs (Figure 7C). The largest MT bundles (34-39 and 40-45 MTs) were found only in PM3, PM4 and metaphase cells (Figure 7D).

**Figure 7.**
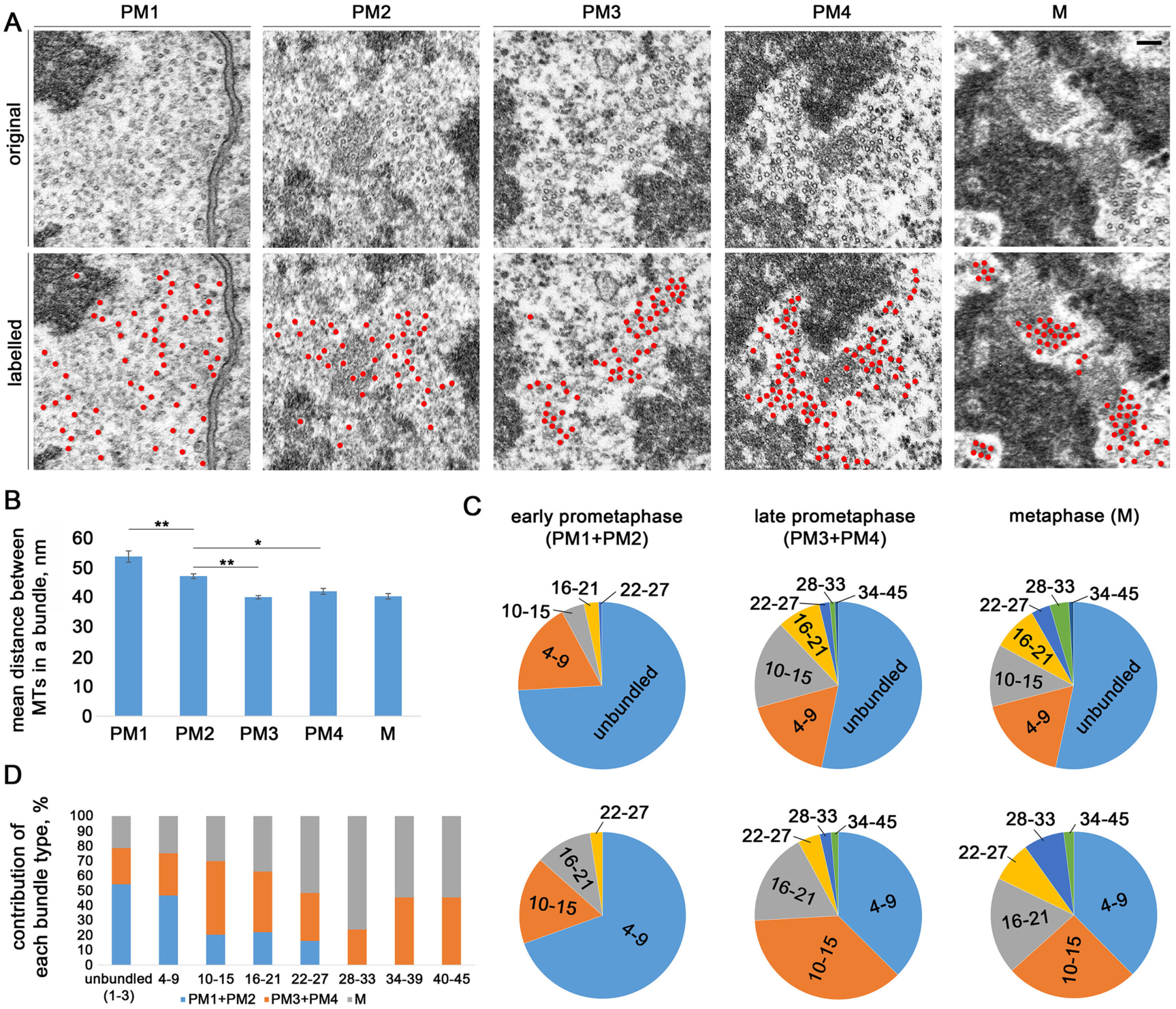
MT organization in S2 cell prometaphases and metaphases. (A) Examples of transverse sections of MTs in prometaphase and metaphase cells; in the bottom panels MTs are pseudocolored in red. In the PM1 cell shown, MTs are not bundled. In the PM2, PM3, PM4 and metaphase (M) cells, MTs are organized in bundles, and the average distance between bundled MTs decreases during progression from early prometaphase to metaphase. Scale bar: 0.1 μm. (B) Quantification of the distances between MTs within bundles; the average distances observed in PM3, PM4 and metaphase (M) cells are significantly smaller than those seen in PM1 and PM2 cells (* and **, p<0.05 and 0.01, respectively; ANOVA with post-hoc Tukey HSD). (C, D) Distributions of MT bundle sizes (number of MTs per bundle) from early prometaphase to metaphase. In (C), the upper row of the pie charts shows the distribution of all MTs, while the lower row show only bundled MTs.

**Figure 8.**
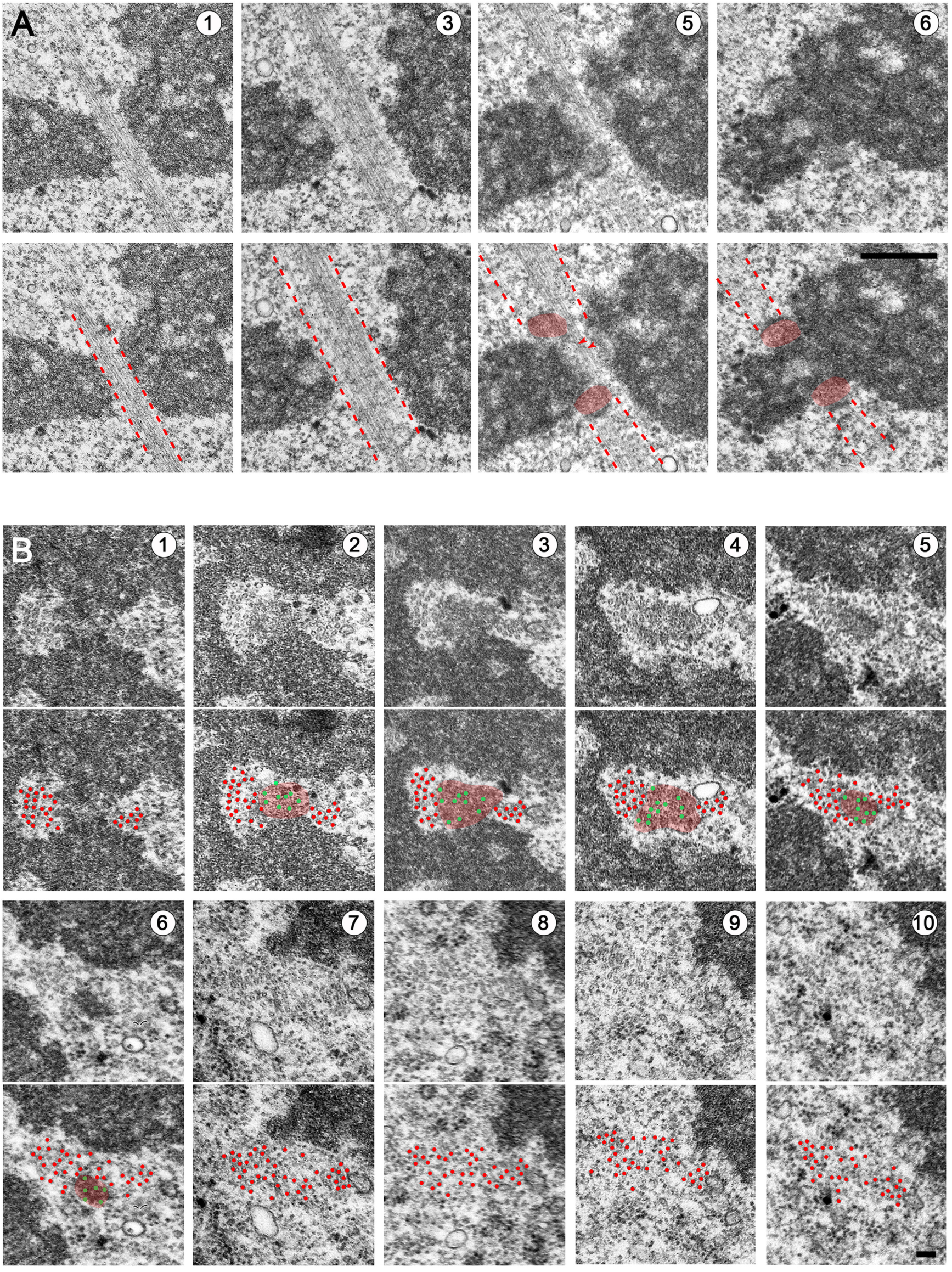
Organization of MT bundles in S2 cell late prometaphase/metaphase. (A) Longitudinal serial sections showing a large MT bundle comprising MTs that are attached to kinetochores (k-fibers) and MTs that run laterally to kinetochores. In the bottom panels, the dotted red lines delimit the MT bundles, and kinetochores are pseudocolored in red. (B) Transverse serial sections through a large MT bundle comprising kinetochore-attached MTs (k-fiber) and two lateral MT bundles. The first and third series of panels show the original images. In the second and forth series of panels, kinetochores and most MTs are pseudocolored in red. MTs embedded into the kinetochore are less clear than the other MTs in the section, and are tentatively pseudocolored in green. Scale bars: 1 μm (A), 0.1 μm (B).

We were not able to visualize mature kinetochore structures before the PM4 stage, in the PM1-PM3 stages the kinetochores are oblong and never show end-on-attached MTs. In PM4 cells, kinetochores have an arched shape similar to that of their metaphase counterparts and often display end-on attached MTs (Figure S4). These observations are consistent with the finding that at mitotic entry human kinetochores appear as large crescents and compact into discrete kinetochore structure only after the attachment to MTs in an end-on fashion (Magidson et al., 2015). Consistent with previous findings (Maiato et al., 2006), longitudinal sections of S2 cell metaphases and early anaphases showed that kinetochores with attached MTs are arched structures that show only an electron dense layer (Figure 3B, 4A, S4, S6A) and fail to exhibit the two typical layers that characterize vertebrate kinetochores. However, after MT depolymerization with colchicine, S2 cell kinetochores exhibit an inner and an outer layer, suggesting that the presence of MTs distorts kinetochore structure (Maiato et al., 2006).

In some transverse sections through metaphase chromosomes, we observed bundles of 9-15 MTs end-on attached to the kinetochores (k-fibers) (Figure 8). Analysis of longitudinal and transverse serial sections (Figure 8A, B) showed that k-fibers contain in average 12 ± 1 MTs, consistent with previous ultrastructural studies on S2 cells (Maiato et al., 2006). We also observed MT bundles running laterally to kinetochores (lateral bundles). In sections including the kinetochore, most lateral bundles contained either 10-14 or 20-30 MTs; in serial sections not including the kinetochores the lateral bundles and the k-bundles formed a single large MT bundle of 30-45 MTs (Figure 8B).

These observations raise the question of the nature of the lateral MT bundles. Are they comprised of antiparallel MTs running from pole to pole with a region of overlap at the cell equator, or are instead MTs that terminate in another kinetochore located proximally or distally with respect to the sectioning plane? Our TEM analyses clearly showed the existence of long pole-to-pole MT bundles (Figure S11) but did not provide sufficient information on the behavior of all lateral MT bundles. To address this question we made mitotic preparation stained for tubulin, Cid and DNA and examined them by a confocal microscope. Cid, the *Drosophila* homologue of CENP-A, is a specific component of centromeric heterochromatin routinely used as a kinetochore marker (Henikoff et al., 2000). Examination of optical sections and threedimensional computer-reconstructed images of late prometaphases and metaphases confirmed the presence of pole-to-pole bundles of approximately the same size as the k-bundles. Examination of particularly favorable preparations revealed that some bundles running adjacent to a kinetochore attach end-on to another kinetochore, while others run till the spindle pole without encountering another kinetochore (Figure 9A, B). We also observed multiple kinetochore/lateral MT bundles converging into larger MT assemblies (Figure 9A, B). Collectively, our confocal and TEM observations are summarized in the 3D model of MT organization during late prometaphase and metaphase (Figure 9 C). This model provides an explanation for the different numbers of MTs in the bundles. Bundles of 9-15 MT are either kinetochore or pole-to-pole bundles, while bundles of 20-30 and 30-45 MTs are likely to comprise different combinations of these two bundle types. The finding that PM3 and PM4 cells exhibit particularly large MT bundles (comprising up to 45 MTs) might reflect the presence of nonaligned prometaphase chromosomes with tandemly arranged kinetochores along the spindle axis.

**Figure 9.**
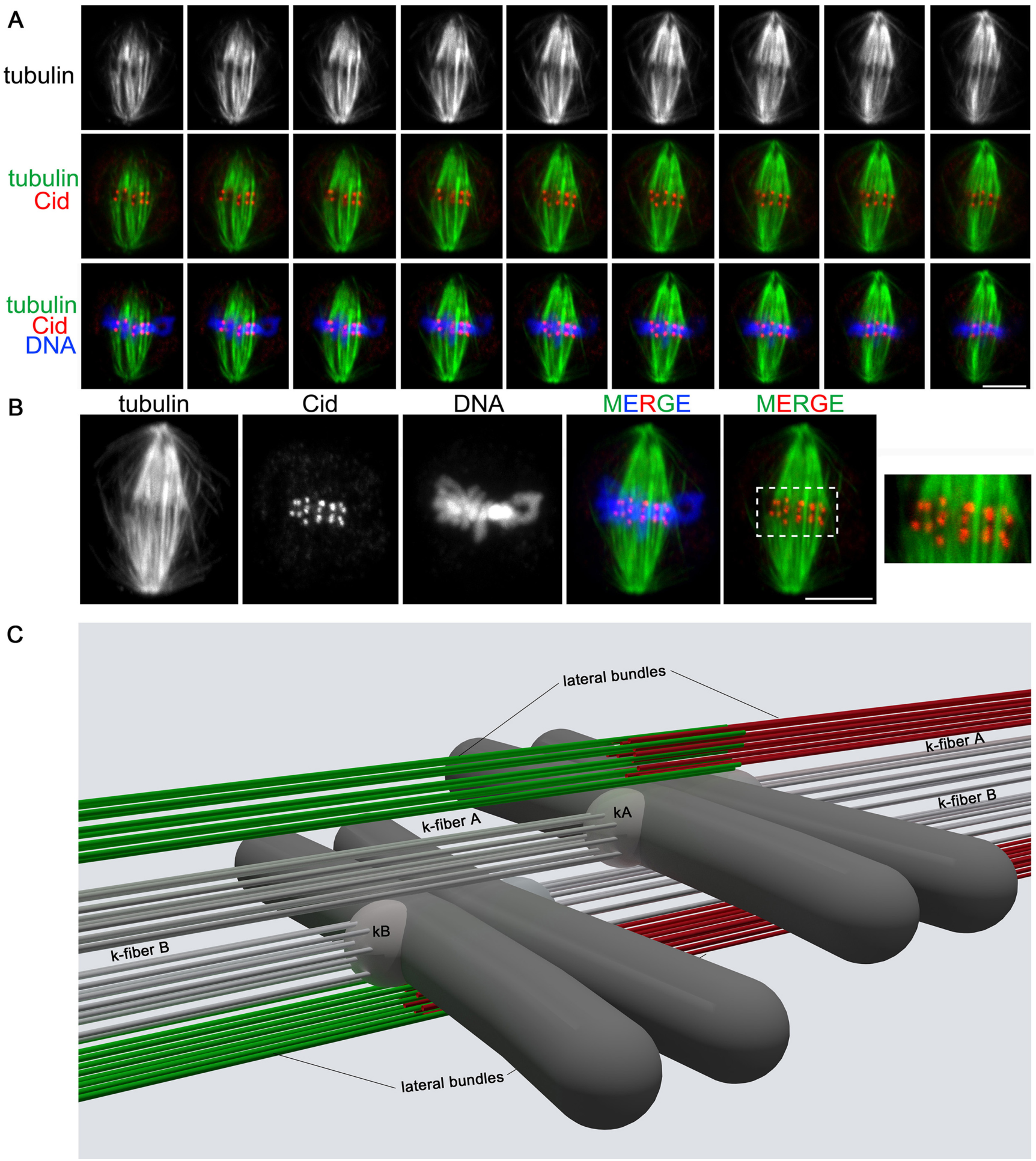
Organization of MT bundles in S2 cell metaphases. (A, B) Serial optical sections (A) and maximum projection (B) of a metaphase cell stained for tubulin, the centromere marker Cid and DNA (DAPI). Some MT bundles are running continuously between the two spindle poles without encountering the centromeres (interpolar bundles), while other bundles end at kinetochores (k-fibers). Note that MT bundles attached to different kinetochores and interpolar bundles are closely apposed (and/or intermingled) in the regions comprised between the centromeres and the spindle poles. Scale bars: 5 μm.(C) A model for MT-kinetochore interaction during late prometaphase and metaphase cells. The model depicts a portion of a late prometaphase/metaphase that includes only two chromosomes. Grey MT bundles are k-fibers; green and red MTs are interpolar MT bundles running laterally to kinetochores; the scheme does not imply that the k-fibers are as numerous as the interpolar MT bundles. k-fibers A and B are attached to the kinetochores of two neighbor chromosomes. Note that k-fiber A is a lateral MT bundle for kinetochore B, while k-fiber B is a lateral MT bundle for the kinetochore A.

The presence of associations between k-fibers of 9-15 MTs and large lateral MT bundles (comprising 9-34 MTs) seems to be peculiar to *Drosophila* S2 cells. MT bundles running laterally to kinetochores have been previously observed by TEM in both longitudinal and cross-sections of several spindle types including those of rat kangaroo PTK cells (Brinkley and Cartwright, 1971; McDonald et al. 1992) and HeLa cells (Wendell et al., 1993; Booth et al., 2011; Nixon et al. 2017), as well as in spindles assembled from *Xenopus* egg extracts (Ohi et al., 2003). However, in all cases, in contrast to S2 cells, the lateral MT bundles appeared to contain much fewer MTs than the kinetochore fibers.

There is evidence that the lateral MT bundles play important roles in chromosome segregation. The most extreme case has been described in the acentrosomal meiotic spindles of *C. elegans* females, where kinetochores are not required for chromosome segregation and chromosomes segregate due to motor protein-mediated interactions with lateral MT bundles (Wignall and Villeneuve, 2009; Dumont et al., 2010; Muscat et al., 2015). In *Drosophila* female meiosis, lateral kinetochore-MT attachments are sufficient for prometaphase chromosome movements but end-on kinetochore-MT attachments are required for homolog bi-orientation, indicating that both lateral and end-on kinetochore-MT interactions cooperate to ensure accurate chromosome segregation (Radford et al., 2015). Also in centrosome-containing spindles of mammalian cells lateral kinetochore-MT interactions have relevant roles in mitotic division. For example, k-fibers of mammalian cells that have lost direct connection to the pole can nonetheless promote chromosome segregation through dynein-mediated lateral interactions with the k-fibers of adjacent chromosomes and/or other non-kinetochore MTs (Sikirzhytski et al., 2014).Consistent with these results, it has been shown that interpolar MT bundles can physically associate with k-fibers forming a bridge between the two sister kinetochores. These bridging MT bundles (called bridging fibers) are thought to balance the forces acting at kinetochores and to contribute to both spindle shaping and chromosome movement (Kajtez et al 2016; reviewed by Tolic, 2017). Based on these studies, we suggest that the peculiar organization of MTs observed in S2 cell metaphase spindle (Figure 9) reflects lateral interactions between kinetochores and both k-fibers and interpolar MTs; these interactions are likely to be functionally important to ensure accurate chromosome segregation

### Concluding remarks

We have shown that S2 cell spindles comprise multiple discrete MT bundles that are likely to interact with each other to ensure proper spindle function. This spindle organization appears to be slightly different from that observed in vertebrate cells.However, we do not know whether it is different from the structure of *Drosophila* embryonic or neuroblast spindles, which are currently incompletely characterized at the ultrastructural level. We have also shown that dividing S2 cells exhibit a peculiar intracellular membrane behavior, which was not previously seen during mitosis of vertebrate cell, *Drosophila* embryonic nuclei, or *Drosophila* larval brain cells.Overall, our observations clearly show that besides their characteristic membrane behavior, S2 cell mitoses are more “open” than their embryonic or neuroblast counterparts. This finding raises the interesting question of whether this feature of S2 cell mitosis is also present in the late (cellularized) embryonic cells from which the S2 lineage derives, or was instead acquired during the process of S2 cell immortalization. Distinguishing between these alternatives and defining whether different *Drosophila* cell types exhibit variations in their mitotic structures will require further ultrastructural studies in additional fly tissues and their derivative cell lines (Lee et al., 2014).

## Materials and methods

### Cell culture and RNA interference

The origin of the *Drosophila* S2 cell line used here has been described previously (Strunov et al., 2016). S2 cells and S2 cells expressing tubulin-GFP were maintained at 25 C in Shields and Sang M3 medium (Sigma) supplemented with 20% heat-inactivated fetal bovine serum (FBS, Invitrogen). The cell density was kept below 6-8×10^6^ cells/ml to avoid formation of cell aggregates. *cnn* dsRNA production and RNAi treatments were carried out according to (Somma et al., 2008); dsRNA-treated S2 cells were grown for 5 days at 25°C, and then processed for TEM analysis.

### Transmission electron microscopy (TEM)

For preparation and ultrastructural analysis of S2 cells we used the protocol described in detail by Strunov et al. (2016). Briefly, the cell pellet was fixed in 2.5% glutaraldehyde in 0.1 M sodium cacodylate buffer for 1 h at room temperature, postfixed for 1 h in 1% water solution of osmium tetroxide containing few crystals of potassium ferricyanide (K_3_[Fe(CN)_6_]), and then incubated overnight at 4°C in 1% aqueous solution of uranyl acetate. Next day, specimens were dehydrated in ethanol series and acetone, and embedded in Agar 100 Resin (Agar Scientific, Essex, UK).Complete polymerization of samples was conducted by keeping them in 60°C oven for three days. Ultra-thin (70 nm) sections were obtained with Leica ultracut ultramicrotome. Sections were examined with JEOL JEM-100SX transmission electron microscope at 60 kV in Inter-institutional Shared Center for Microscopic Analysis of Biological Objects (Institute of Cytology and Genetics, Novosibirsk, Russia). The obtained images were slightly modified in Photoshop CS5 using levels and brightness and contrast tools. 3D reconstructions of prometaphase cells were obtained using the “Reconstruct” free editor (Fiala, 2005).

### Measurements with ImageJ

The length of membranes (NME, RDM, QNM and ER) in S2 cells at different mitotic stages was measured with ImageJ software (https://imagej.nih.gov/ij/), using the simple freehand selection tool and measure function. For membrane measurements, we considered only longitudinal or slightly oblique longitudinal sections. The data obtained for each mitotic stage were compared using one-way ANOVA with post-hoc Tukey HSD test.

The distance between MTs within a bundle at different mitotic stages was also measured using ImageJ software. For each mitotic stage, we analyzed 10 transversally cut bundles containing at least 10 MTs. For each bundle, we measured the radial distances between the MT at the center of the bundle and its neighbor MTs; measures were taken from MT center to MT center and the mean distance was quantified. Data obtained for different mitotic stages were compared using one-way ANOVA with post-hoc Tukey HSD test.

### Confocal microscopy observations

2× 10^6^ of S2 cells were centrifuged at 200 × *g* for 5 min, washed in 2 ml of phosphate buffered saline (PBS, Sigma), and fixed for 10 min in 2 ml of 3.7% formaldehyde in PBS. Fixed cells were spun down by centrifugation at 200 × *g* for 5 min, resuspended in 500 μl of PBS and placed onto a clean slide using Cytospin™ 4 cytocentrifuge (Thermo Fisher Scientific) at 900 rpm for 4 min. The slides were then immersed in liquid nitrogen, washed in PBS, incubated in PBT (PBS + 0.1% TritonX-100) for 30 min and then in PBS containing 3% bovine serum albumin (BSA) for 30 min. The following primary antibodies were used for immunostainings (all diluted in PBT): mouse monoclonal anti-α-tubulin (1:600, Sigma, T6199), mouse monoclonal anti-Lamin Dm0 (1:300, Developmental Studies Hybridoma Bank, ADL67.10), rabbit polyclonal anti-Cid (1:300, Abcam, ab10887) and rabbit polyclonal anti-GFP (1:200, Invitrogen, A11122). These primary antibodies were detected by incubation for 1 h with FITC-conjugated anti-mouse IgG (1:30, Sigma, F8264), Alexa Fluor 568 conjugated anti-mouse IgG (1:500, Invitrogen, A11031), Alexa Fluor 488 conjugated anti-rabbit IgG (1:300, Invitrogen, A11034) and Alexa Fluor 568 conjugated antirabbit IgG (1:350, Invitrogen, A11077). All slides were mounted in Vectashield medium with DAPI (Vector Laboratories, H-120) to stain DNA and reduce fluorescence fading. Confocal immunofluorescent images were obtained on a Zeiss LSM 710 confocal microscope, using an oil immersion 100X/1.40 plan-apo objective and the ZEN 2012 software.

## Acknowledgements

We thank Alexey Khodjakov for a critical reading of the manuscript and helpful suggestions. This work was supported by a grant from the Ministry of Education and Science of Russian Federation (14.Z50.31.0005), by a Russian Science Foundation project (16-14-10288) for confocal analysis of lamin behavior, and by Fundamental Scientific Research Programs (0324-2016-0003 and 0310-2016-0005) for TEM analysis and RNAi, respectively.

## List of abbreviations

DNM: double nuclear membrane
ER: endoplasmic reticulum
MT: microtubule
NEBD: nuclear envelope breakdown
NPC: nuclear pore complex
QNM: quadruple nuclear membrane
RDM: residual double membrane
SE: spindle envelope
TEM: transmission electron microscopy

